# Shared symmetry detection with divergent temporal dynamics in marmosets and humans

**DOI:** 10.64898/2026.08.17.745359

**Authors:** Amirhossein Asadian, Dirk B. Walther, Peter J. Kohler, Liya Ma

## Abstract

Which features of human perception are shared across the primate lineage, and which are specializations, is a central question in the biology of cognition. fMRI studies have located symmetry-selective cortex in macaques [1,2], and preliminary intracortical recordings point to symmetry-sensitive neurons in V2–V4 [3]. Yet no study has measured symmetry responses in non-human primates (NHPs) with the same non-invasive electrophysiology used in humans. Thus, it remains unknown whether NHPs respond to symmetry with dynamics that match those in humans. Using steady-state visual evoked potentials during passive viewing, we compared responses to two well-matched wallpaper groups, double reflection (PMM) and four-fold rotation (P4), in common marmosets and humans, using identical stimuli. Marmosets showed robust reflection-symmetry responses comparable in relative magnitude, temporal dynamics and scalp distribution to those of humans. Rotation responses were also reliable but transient, without the sustained late component prominent in humans. This dissociation suggests that fast, feedforward computations underlying symmetry detection are conserved across primates, whereas sustained processing that elaborates symmetry representations, likely recurrent or feedback in origin, is reduced in marmosets and most clearly for rotation. These findings identify a point of divergence for primate visual processing and are consistent with the emerging view that marmosets rely less on cortical feedback than macaques or humans. Our approach establishes marmosets as a tractable model for symmetry processing and opens a path to intracortical recording in easily accessible extrastriate cortex.

## 1. Introduction

Symmetry is a pervasive feature of natural scenes, biological forms, and human artifacts and a fundamental organizing principle of mid-level vision [4,5]. While many species are sensitive to visual symmetry [6], the brain mechanisms that drive these responses remain understudied outside of humans, including non-human primates. The current study aimed to investigate whether human symmetry processing reflects computations shared across primates or includes specializations more developed in humans.

Reflection symmetry has long been considered privileged relative to other two-dimensional regularities: It is detected more rapidly [7,8] and is more salient [7,9,10]. Reflection is also the most prevalent symmetry type in the body shapes of virtually all mobile animals and therefore a particularly relevant cue for face, body, and animacy perception. Rotation symmetry preserves appearance under rotation rather than mirror reflection and is less readily associated with evolutionary pressures. Reflection elicits stronger neural responses than rotation symmetry [11], but the human visual system responds robustly and systematically to both reflection and rotation, and represents 2D regularities hierarchically [12]. Regular textures containing double reflection (PMM) and four-fold rotation (P4) thus provide a well-controlled test case for whether different symmetry types are processed similarly across primate species.

In humans, the neural basis of symmetry perception has been reliably localized to a specific network of extrastriate regions [1]. Along the visual processing hierarchy, systematic responses to symmetry first emerge in area V3 and produce the strongest responses in areas V4, VO1 and Lateral Occipital Complex (LOC) [13]. Electrophysiological work has further characterized the temporal dynamics. Event-related potential (ERP) studies have identified a Sustained Posterior Negativity (SPN) emerging 200–250 milliseconds after stimulus onset in electrodes over occipital cortex, that scales parametrically with symmetry salience [11]. Steady-State Visual Evoked Potentials (SSVEPs) paradigms provide an isolated, high signal-to-noise ratio read-out of symmetry-specific responses in the odd harmonics of the stimulation frequency and reveal temporal dynamics that are consistent with the SPN [12,13].

Recent studies with macaque monkeys provide evidence for a homologous network for symmetry processing in non-human primates: Functional MRI has revealed that symmetry responses are highly consistent between macaques and humans, both in terms of patterns of responses to individual symmetry types and brain areas involved [2]; notably, macaque V3 and V4 already show a reflection-over-rotation gradient of roughly one third, mirroring the same gradient documented in humans [2,12]. Intracranial recordings in macaque IT have further suggested that symmetry preference may arise from generic computations of object distinctiveness [14]. Efforts to extend this work with electrophysiological recordings within V3 and V4 are impeded because these areas are buried within the lunate sulcus in macaques, presenting challenges especially for the laminar recordings necessary to dissect local circuit mechanisms [15]. The common marmoset (Callithrix jacchus) provides a promising alternative. This New World primate combines a high-acuity visual system homologous to that of humans with a largely lissencephalic cortex that exposes V2, V3, V4, and V3A on the cortical surface [15,16], enabling chronic, high-density laminar recordings from regions inaccessible in macaques. Recent work in comparative connectomics has identified another difference between marmoset and macaque cortex that may have direct bearing on symmetry processing: marmoset cortex is on average more feedforward than macaque cortex, and strong top-down projections from prefrontal to posterior parietal cortex seen in macaques are weak or absent in marmosets [17]. If feedback and recurrent processing are reduced in the marmoset, symmetry computations that depend on them should be affected more than those that do not. This yields a concrete cross-species prediction that can be tested using SSVEP measurements at the scalp.

The utility of marmosets as a model for studying mid-level vision depends critically on the presence of shared mechanisms between humans and marmosets. Here we present the first assessment of this, and to our knowledge, the first measurement of any kind of brain responses to symmetry in marmosets. We collect scalp EEG with a SSVEPs design that allows us to compare symmetry responses between marmosets and human participants. Reflection symmetry responses were comparable in magnitude, temporal dynamics and topography across the two species. Rotation symmetry also produced measurable responses in marmosets, but they lacked the well-documented sustained component seen in humans. These findings identify the feedforward computations that support symmetry detection as candidate primate-general mechanisms and establish the marmoset as a tractable model for their mechanistic investigation, while highlighting a specific late-stage difference between species that motivates targeted follow-up.

## 2. Materials and Methods

### 2.1. Animal subjects

All procedures were conducted on three naïve adult male common marmosets (*Callithrix jacchus* subjects F, M, and T; all 4 years of age; weights 400, 450, and 460 g, respectively). All experimental procedures adhered to the guidelines of the Canadian Council on Animal Care and were approved by the Animal Care Committee at York University.

### 2.2. Human participants

13 participants (7 females, mean age 25.7 ± 8.0) took part in the EEG experiment. All participants were pre-screened to confirm that they had normal or corrected-to-normal visual acuity on the Bailey-Lovie chart, and no neurological conditions or history of head injury. Informed consent was obtained before the experiment, and all experimental procedures were conducted as specified in a protocol approved by the Office of Research Ethics at York University.

### 2.3. Marmoset EEG grid and chamber design

For each animal, a flexible EEG grid and an individualized recording chamber (Figure 1B) were surgically implanted under anaesthesia, using aseptic techniques. Individualized recording chambers were designed based on each animal’s skull model, constructed from averaged T1-weighted MRI scans, and were covered with a custom-designed cap secured using set screws (Figure 1B). Both the chamber and cap were 3D-printed, with a combined weight of 6 g. The 32-channel flexible grid was custom designed with 7 rows spanning the dorsal skull surface (Figure 1C). The rows were spaced by 4.5 mm, and contacts within a row were spaced by 3 mm. They were manufactured by NeuroNexus Technologies (Ann Arbor, MI, USA). From posterior to anterior, the rows were named O, PO, P, PC, C, F, AF (Figure 1C), with the central sites located approximately above V1, border of V1/V2, border of V2/V6, border of V6/IPS areas, area PE, area 6M, and area 9, respectively.

**Figure 1.**
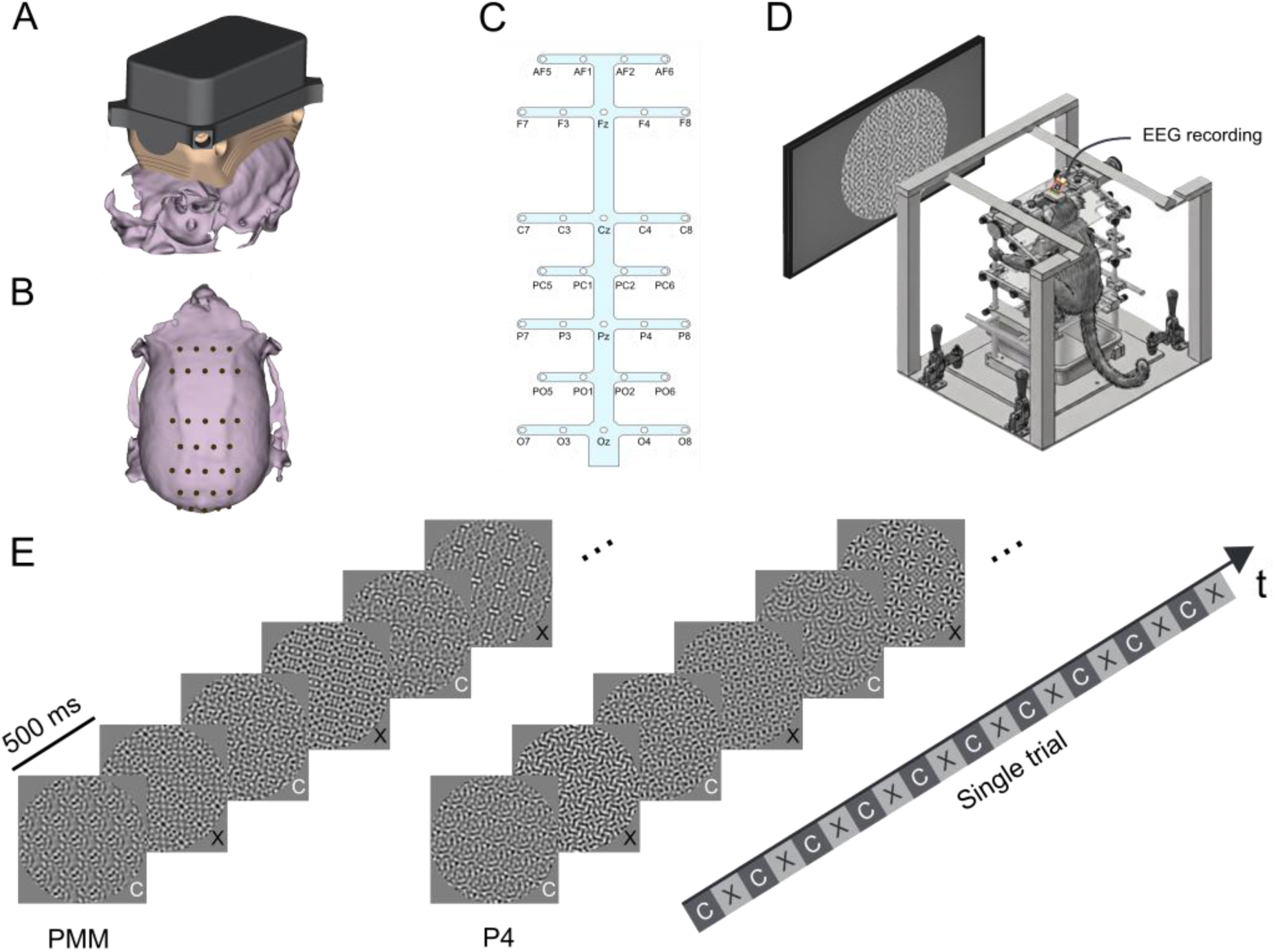
Experimental setup, EEG configuration, and behavioural task. A, Customized chamber and protective cap positioned on the skull. B, Recording site target locations with reference to the skull. C. Flexible EEG grid layout. D, Marmoset training apparatus during visual stimulation for steady-state visually evoked potential (SSVEP) recording. E, Behavioural task schematic illustrating alternating presentation of control exemplars (C) and PX exemplars (PMM or P4 image groups). Each trial consists of 10 image pairs, with each image presented for 500 milliseconds.

### 2.4. Marmoset surgical procedure

Following acclimation to the marmoset chair (Neuronitek, London, ON, Canada), animals underwent an aseptic stereotaxic surgery to implant the EEG grid and head chamber. Sedation was induced with ketamine (20mg/kg); and anesthesia was maintained with 2% isoflurane. Depth of anesthesia was monitored throughout the procedure. A midline scalp incision was made, the temporalis muscles were retracted to expose the cranium, and the periosteum and residual connective tissue were removed. For the EEG ground, a titanium screw was placed above the right somatosensory cortex (approximate coordinates: AP: +7mm; ML: +5mm), and the ground wire was tightly wound around it. The grid was positioned such that the PCz site was placed at the coordinates of AP = −0.4mm from interaural zero, and ML = 0. Drops of sterile saline were applied to the grid, so it adhered to the skull. Thin strips of dental cement (Duo-Link Universal; Bisco, Schaumburg, IL, USA) were then applied conservatively over the grid to secure it in position. Following grid placement, the skull surface within the intended chamber margin was mechanically roughened using a small stainless-steel brush, thoroughly rinsed, and dried. Two coats of dental adhesive resin (All-Bond Universal; Bisco) were applied, air-dried, and cured with ultraviolet light. More dental cement was applied to secure the chamber at its predetermined location, thoroughly cover the grid and ground screw within the chamber and create a fluid-tight seal around it. The ground and reference wires from the grid were then securely connected with the ground wire. Finally, a dust cap was secured over the chamber with set screws to protect the connector and wires.

### 2.5. Marmoset apparatus and data acquisition

After recovery, the animals were acclimated to being head-fixed by the head chamber with a Kopf cube head restraint (Neuronitek) (Figure 1A). The setup was adjusted so the marmosets viewed the centre of an LCD monitor (BenQ XL2420T; 120 Hz refresh rate) from a 57-cm distance. Eye position was monitored continuously via infrared tracking of the pupil using a binocular eye tracker (TrackPixx3, 2 kHz; VPixx Technologies, Montreal, QC, Canada). EEG signals were acquired at 30KHz using a multichannel recording system (RHS stim/recording controller, Intan Technologies, Los Angeles, CA, USA) via a 32-channel headstage. Continuous EEG was acquired at 30 kHz, imported from the Intan system, and reordered to match the physical 32-contact grid layout spanning occipital to frontal cortex. Eye-movement data were acquired synchronously and sent to the recording system via a Datapixx3 interface (VPixx Technologies). Visual stimuli were presented on the monitor through Datapixx3, and the behavioural task was controlled by custom MATLAB (2022b; MathWorks, Natick, MA, USA) scripts using Psychtoolbox (version 3.0.18). To ensure precise measurement of visual stimulus onset, an in-house built photodiode circuit was affixed to the monitor to detect the exact timing of each image onset on each trial. Trials in which photodiode signals indicated imprecise or ambiguous onset timing were excluded from further analysis (see section *2.7*. *Data analysis*). After each completed trial, animals received a marshmallow juice reward delivered via a peristaltic pump (Adafruit, Brooklyn, NY, USA).

### 2.6. Human data acquisition

Human data were acquired using a Magstim EGI EEG system consisting of 128-channel HydroCell Geodesic Sensor Nets, a Net Amps 400 amplifier and Netstation software version 5.5 (Magstim EGI, Eugene, OR). Stimuli were presented on a Dell P2417h monitor running 1920 ✕ 1080 resolution at 60 Hz. Stimulus presentation and synchronization with EEG acquisition was done using in-house software. Continuous EEG data initially acquired at 500 Hz was resampled at 420 Hz to provide seven samples per video frame.

### 2.7. Stimulus generation

Visual stimuli were patterned textures drawn from two wallpaper symmetry groups, P4 and PMM, together with corresponding control images. Exemplars and controls were generated following the procedures described by Kohler et al. (2016), with adaptations for presentation in marmosets. Exemplars for each group were created from random-noise textures transformed according to the symmetry operations defining P4 (fourfold rotation symmetry) or PMM (double reflection symmetry). All stimuli were rendered as grayscale images and presented within a circular aperture subtending approximately 24° of visual angle at a viewing distance of 57 cm. Because animals were not required to maintain central fixation, a larger stimulus size compared to prior work was used so that effective stimulation of the animal’s central visual field was less dependent on precise fixations. For each symmetry group, 24 unique exemplars were generated to minimize adaptation to specific low-level image features across trials. Each exemplar was paired with a matched control image that preserved the original Fourier amplitude spectrum while disrupting its higher-order symmetry structure, using the same phase-scrambling procedure as prior work. The phase-scrambling preserves the tiling of the original wallpaper group patterns, and therefore produces another wallpaper group, P1, that only contains translation symmetry. Contrast and mean luminance were matched across all stimuli, which were presented on a mid-grey background. Because animals were not required to maintain strict fixation, stimuli were made large enough to ensure that symmetry structure remained present at the fovea regardless of small eye movements during the trial.

### 2.8. Experimental Design

We used a Steady-State Visual Evoked Potential (SSVEP) design where each trial consisted of a sequence of control and symmetry images. In Steady-State Visual Evoked Potentials paradigms periodic stimulation gives rise to narrow-band peaks in the response spectrum at integer multiples of the stimulation frequencies, called harmonics [18]. In our design, a stimulus cycle consisted of a control image followed by a symmetry image, each showed for 500 ms. On each trial, 10 stimulus cycles were presented, with 10 unique control-symmetry image pairs (Fig. 1C). By definition, even harmonics capture the component of the response that is identical for the first and second half of the cycle, whereas odd harmonics captures the component of the response that differs between the first and second half. This means that in our design, the even harmonics will capture low-level visual responses that are the same for symmetry and control images, while the odd harmonics will capture the symmetry-specific response [13].

For marmoset data, from a total pool of 24 exemplars per symmetry group, three fixed subsets of 10 were used across sessions (exemplars 1–10, 11–20, and 13–22), with the number of sessions using each subset varying across animals (Marmoset F: 1, 6, 3 sessions respectively; Marmoset M: 7, 4, 1; Marmoset T: 3, 1, 3). P4 and PMM trials were intermixed and presented in a pseudorandom order across the session. The visual system is known to respond more strongly to horizontal and especially vertical axes of reflection symmetries [7,19]. To avoid this cardinal axis bias and reduce adaptation to specific retinal configurations, the axis of symmetry for each image pair was drawn from a discrete set of oblique orientations (15°, 45°, 75°, 105°, 135°, 165°), thereby explicitly avoiding horizontal (0°/180°) and vertical (90°) axes. The human data were acquired using the same stimuli, with two minor modifications: First, symmetry axes were not obliquely oriented. Second, a subset of 10 of the 24 exemplars was selected for each condition, and used across all trials. The number of trials was variable among the human participants, with most participants doing 25 or 30 trials per condition, and some doing as many as 50.

### 2.9. Marmoset training procedures

Following a two-week postoperative recovery period after EEG grid implantation, animals began head-fixed training to habituate them to the restraint apparatus and head fixation. Once animals tolerated head fixation, we initiated fixation training for oculomotor calibration by presenting marmoset face images as fixation targets; correct fixations were rewarded with marshmallow juice. At the start of each recording session, a brief fixation block was used to perform eye-tracker calibration, after which animals began completing the SSVEP trials. To ensure cross-species comparability, the SSVEP trial structure was matched to the human paradigm: each trial consisted of 10 image pairs presented over a 10-second duration. This structure was introduced incrementally; as the animals’ tolerance improved, session length was progressively increased from 50 to 100 trials per session.

### 2.10. Data analysis

For both marmosets and humans, signals were demeaned on a per-channel basis and notch-filtered at 60 Hz and 120 Hz using second-order IIR notch filters to attenuate line noise and its first harmonic. To prevent aliasing before down sampling, we applied a zero-phase finite-impulse-response (FIR) low-pass filter (cutoff 400 Hz) and then resampled the data to 1 kHz, after which a zero-phase 0.1–30 Hz band-pass FIR filter was used to isolate the low-frequency components carrying the 1 Hz SSVEP and its low harmonics. The same preprocessing pipeline was applied to all channels; no additional re-referencing was performed beyond the recording system’s reference configuration. Eye-position analog signals were low-pass filtered at 400 Hz and down sampled to 1 kHz for subsequent quality control.

Since the EEG grid was chronically fixed to the skull, data from all recording sessions were pooled within each animal and analysed at the single-subject level. Because calibration quality and eye stability varied across sessions, we applied additional inclusion criteria based on raw horizontal eye-position traces. For each trial, we extracted horizontal eye position over the full image sequence, identified signal-loss events (for example, eye closures) longer than 500 ms, and computed session-specific global eye position thresholds (mean ± 4 SD) across all trials, for each session. Thresholds were validated by comparing them to the distributions observed in sessions with high-quality calibration. Time-series data were rejected on a epoch basis, where an epoch consisted of a single image pair. If the horizontal position exceeded the thresholds or overlapped with long signal-loss events, the entire epoch was rejected. Trials with more than three rejected epochs (out of a total of 10) were excluded from further analysis.

For each animal (F, M, T), we applied rigorous exclusion criteria to ensure data quality. Trials were included only if they met these conditions: (i) all image pairs in the trial passed behavioural, photodiode-based, and eye-based quality checks and (ii) the trial contained the full 10 image pairs after quality checks. This strict selection resulted in the following final datasets: Animal F contributed 10 sessions (P4: 265/470 trials; PMM: 272/470 trials); Animal M contributed 13 sessions (P4: 399/639 trials; PMM: 355/621 trials); and Animal T contributed 5 sessions (P4: 151/348 trials; PMM: 150/332 trials).

From each accepted trial, we extracted an 8-s EEG segment starting at the onset of the third image in the sequence, treating the first and last alternation cycles as prelude and postlude periods and excluding them to minimize onset/offset transients in the steady-state response. This 8-s segment (8 control–symmetry alternations at 1 Hz) was partitioned into four non-overlapping 2-s blocks, which were averaged to yield a single 2-s waveform per trial, channel, and condition (P4, PMM). Finally, a Discrete Fourier Transform (DFT) was computed on this averaged waveform to obtain the full complex frequency spectrum for each epoch.

To visualize the time course of the SSVEP responses, we reconstructed waveforms from selected harmonic subsets using the ROI-level complex spectra. For ROI analyses, the marmoset occipital ROI comprised channels O3, Oz, and O4, corresponding to the custom 32-channel grid; the human occipital ROI comprised channels 70, 74, 75, 81, 82, and 83 from the 128-channel montage. For a given harmonic set (odd: 1, 3, 5, … 19 Hz; even: 2, 4, 6, … 20 Hz, yielding 10 harmonics per set) we created a frequency-domain vector of length N, inserted the appropriately rescaled complex coefficients at the bins corresponding to the chosen harmonics, imposed conjugate symmetry, and applied the inverse FFT to obtain 1-s, 1-kHz time-domain waveforms. For marmoset data, waveforms were reconstructed trial-by-trial for each condition (P4, PMM), and mean ± SEM across trials are shown. For human data, waveforms were reconstructed trial-by-trial per subject, coherently averaged across trials within each subject, and then incoherently averaged across subjects (mean ± SEM across subjects). For topographic analyses, complex coefficients at specific harmonics (e.g., 1 Hz, 2 Hz, or pooled odd/even sets) were averaged across trials within each animal and across subjects for human data. Amplitudes were computed as the absolute value of the coherent mean complex coefficient at each electrode. For marmoset data, electrodes were mapped onto a 2D representation of the custom 32-channel grid; for human data, 3D electrode positions were projected onto 2D using azimuthal equidistant projection. In both cases, amplitude values were spatially interpolated onto a regular grid using Delaunay triangulation and natural-neighbour interpolation to generate smooth scalp maps. All preprocessing and analyses were implemented in MATLAB (R2022b; MathWorks, Natick, MA) using custom scripts.

### 2.11. Statistical analysis

All statistical tests were conducted separately for each animal. To assess whether SSVEP responses at a given harmonic were reliably different from zero, we treated the complex Fourier coefficients across trials as bivariate observations. For marmosets, we applied the one-sample circular T² statistic [20] against the null hypothesis of zero mean complex response, separately for each harmonic, condition, and electrode. For humans, we applied one-sample Hotelling’s T² tests on the real and imaginary parts of the complex coefficients against the same null. Given the large number of trials per condition, the complex Fourier coefficients are approximately bivariate-normally distributed across trials. Multiple comparisons across harmonics and electrodes were controlled using false-discovery-rate correction with a threshold of α = 0.01; electrodes surviving this correction are indicated in the figures. To compare symmetry conditions, we tested differences between PMM and P4 using a two-sample circular T² statistic [20] on the complex Fourier coefficients at the harmonic of interest (e.g., 1 Hz, 2 Hz) for marmosets. For humans, symmetry conditions were compared at the subject level using paired Hotelling’s T² tests on the ROI-averaged complex coefficients, with FDR correction across harmonics. For ROI analyses, the same procedures were applied to the ROI-averaged complex amplitudes. All reported effects thus reflect differences in both amplitude and phase of the complex SSVEP response, evaluated within a unified multivariate framework. All analyses were implemented in MATLAB (R2022b; MathWorks, Natick, MA) using custom scripts. For human paired Hotelling’s T² tests, multivariate effect size is reported as Mahalanobis *D*.

## 3. Results

Three adult male marmosets contributed 28 recording sessions (F: 10; M: 13; T: 5). After quality control, the final marmoset dataset comprised 815 P4 and 777 PMM trials. The human sample comprised 13 participants (7 female; mean age 25.7 ± 8.0years), who completed 25–50 trials per condition.

To determine whether PMM and P4 elicited reliable SSVEP responses in marmosets and humans, and to characterize condition differences across response components associated with image alternation and symmetry processing, we quantified response amplitudes in an occipital region of interest at the first three even harmonics (2, 4, and 6 Hz) and first three odd harmonics (1, 3, and 5 Hz). The choice to focus on the first three harmonics was based on previous work using a similar paradigm in humans, which found that symmetry-specific responses were most pronounced in the first three odd harmonics [13]. Both even and odd harmonics produced robust responses in all three marmosets and in our group-level analysis of the human participants (Figure 2). Among the even harmonics, marmoset F showed no P4–PMM difference at 2 Hz (two-sample circular T² = 0.021, n_P4_ = 265, n_PMM_ = 272, p_FDR_ = .175), 4 Hz (T² = 0.003, p_FDR_ = .828), or 6 Hz (T² = 0.001, p_FDR_ = .828). Marmoset M showed significantly larger P4 than PMM responses at 2 Hz (two-sample circular T² = 0.161, n_P4_ = 399, n_PMM_ = 355, p_FDR_ < .001), 4 Hz (T² = 0.141, p_FDR_ < .001), and 6 Hz (T² = 0.040, p_FDR_ < .001). Marmoset T also showed no even-harmonic differences at 2 Hz (two-sample circular T² = 0.055, n_P4_ = 151, n_PMM_ = 150, p_FDR_ = .051), 4 Hz (T² = 0.032, p_FDR_ = .122), or 6 Hz (T² = 0.028, p_FDR_ = .122). In the human group, even-harmonic amplitudes differed modestly between conditions, with PMM greater than P4 at 2 Hz (paired Hotelling’s T² (2, 11) = 13.65, Mahalanobis D = 0.339, p_FDR_ = .023) and 4 Hz (T² (2, 11) = 9.64, D = 0.254, p_FDR_ = .039), and P4 greater than PMM at 6 Hz (T² (2, 11) = 39.02, D = 0.234, p_FDR_ = .001). By contrast, odd harmonics showed the clearest condition effect at 1 Hz: PMM exceeded P4 in all three marmosets—F (T² = 0.183, n_P4_ = 265, n_PMM_ = 272, p_FDR_ < .001), M (T² = 0.531, n_P4_ = 399, n_PMM_ = 355, p_FDR_ < .001), and T (T² = 0.206, n_P4_ = 151, n_PMM_ = 150, p_FDR_ < .001)—and in the human group (paired Hotelling’s T² (2, 11) = 19.69, Mahalanobis D = 0.825, p_FDR_ = .014). At 3 and 5 Hz, marmosets showed mixed condition effects, whereas the human group showed no condition difference at 3 Hz (paired Hotelling’s T² (2, 11) = 2.84, D = 0.325, p_FDR_ = .466) or 5 Hz (T² (2, 11) = 0.98, D = 0.208, p_FDR_ = .649). Overall, the pattern is largely consistent across marmosets and humans, with the strongest and most reliable condition difference observed at 1 Hz and more variable effects at the higher harmonics.

**Figure 2.**
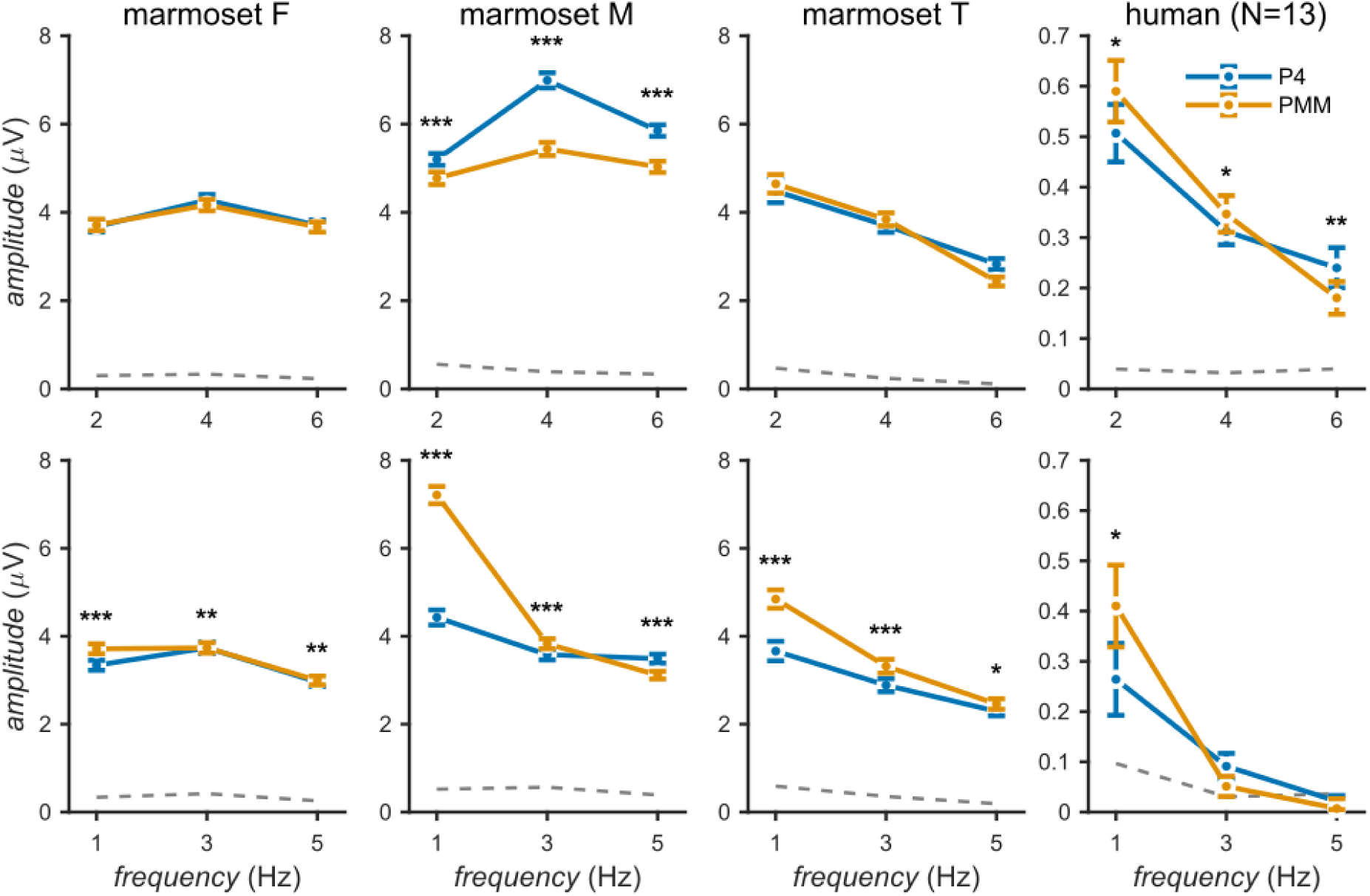
ROI harmonic amplitudes for even and odd frequency components across marmosets and humans. Mean ± SEM response amplitudes are shown at the first three even (2, 4, 6 Hz; top row) and first three odd (1, 3, 5 Hz; bottom row) harmonics of the 1 Hz stimulation frequency, for P4 (blue) and PMM (red) conditions. Amplitudes were computed as the magnitude of the mean complex Fourier coefficient averaged across ROI channels (marmosets: Oz, O3, O4; humans: six electrodes over occipital cortex). For marmosets, values reflect the mean ± SEM across trials; for humans, values reflect the mean amplitude across subjects with asymmetric error bars given by a 2D error ellipse in the complex plane. The grey dashed line indicates the estimated noise floor, computed as the coherent average amplitude within each noise frequency band per condition, then averaged incoherently across bands and conditions. Asterisks indicate FDR-corrected condition differences at each harmonic (\**: p < 0.05, **: p < 0.01**, \*\*\****: p < 0.001; marmosets: two-sample circular T² test on complex coefficients; humans: paired Hotelling’s T² test on complex coefficients). Columns correspond to individual marmosets (F, M, T) and the human group.

To investigate the temporal dynamics of the responses, we reconstructed single-cycle average waveforms by selectively retaining harmonic subsets in the Fourier domain and transforming them back into the time omain. When only the even harmonics of the 1 Hz stimulation frequency were retained, the resulting waveform captures transient responses time-locked to each image alternation, reflecting the visual system’s response to the change in image content at each 500 ms transition. These even-harmonic waveforms were highly similar in shape between the P4 and PMM conditions across all animals and in the human group, indicating that the low-level transient response to image alternation did not differ between symmetry types (Figure 2, middle row). In contrast, reconstructions retaining only the odd harmonics revealed a sustained response that persisted throughout each 500 ms half-cycle rather than returning to baseline between image alternations. In humans, both P4 and PMM elicited sustained odd-harmonic responses of comparable duration, with PMM showing a larger amplitude. In marmosets, the PMM condition similarly produced a sustained deflection, whereas the P4 response was more transient, returning closer to baseline within the half-cycle; this difference in response persistence between conditions was consistent across animals, with PMM also showing larger amplitude in two of three marmosets (Figure 2, bottom row). The P4 and PMM conditions produced waveforms of similar overall morphology in the raw EEG (Figure 2, top row), with differences emerging specifically in the odd-harmonic reconstruction.

To characterize the spatial distribution of the even harmonics, we plotted the second harmonic (2f) across the full electrode array. Both PMM and P4 elicited strong 2f responses over posterior electrodes, with a broadly bilateral parietal-occipital maximum (Figure 4). One-sample circular T² tests identified widespread posterior clusters of non-zero even-harmonic responses in each marmoset after false-discovery-rate correction (q < .01). Across significant marmoset electrode-by-animal observations, circular T² values ranged from 0.014 to 2.637 (p_FDR_ ≤ .007). In the human group, one-sample Hotelling’s T² tests likewise identified widespread significant posterior responses (T² (2, 11) = 17.74–120.00, p_FDR_ ≤ .010, Mahalanobis D = 1.168–3.038). Together, these results suggest that our image pairs were well matched in terms of low-level visual properties, both within and between conditions, and that even-harmonic SSVEPs primarily reflect non-specific visual transients driven by image alternation. Together, these results suggest that our image pairs were well-matched in terms of low-level visual properties, both within and between conditions, and that even-harmonic SSVEPs primarily reflect non-specific visual transients driven by image alternation.

The symmetry-specific signal was carried by the odd harmonics and was independent of the image-update response indexed by the even harmonics. Odd-and even-harmonic amplitudes were uncorrelated across animals and conditions (all p > 0.05), and the condition effect at 1f₁ remained significant after partialling out even-harmonic amplitude on each trial (all p < 0.001). The symmetry effect therefore does not reflect a global gain change in response gain shared with the image-update response.

Although the even harmonics primarily index the non-specific image-update transient, they were not fully condition-invariant. Marmoset M showed larger even-harmonic responses for P4 than PMM at all three harmonics (all p < 0.001), whereas the human group showed a smaller, mixed pattern (PMM > P4 at 2 and 4 Hz; P4 > PMM at 6 Hz), and marmosets F and T showed no reliable even-harmonic condition difference. The even-harmonic waveform shape was closely matched between conditions in every case (**Figure 3**), indicating that these were differences in response gain rather than in the form of the image-update response. The inconsistent direction across subjects and species argues against a systematic low-level stimulus difference between the two wallpaper groups.

**Figure 3.**
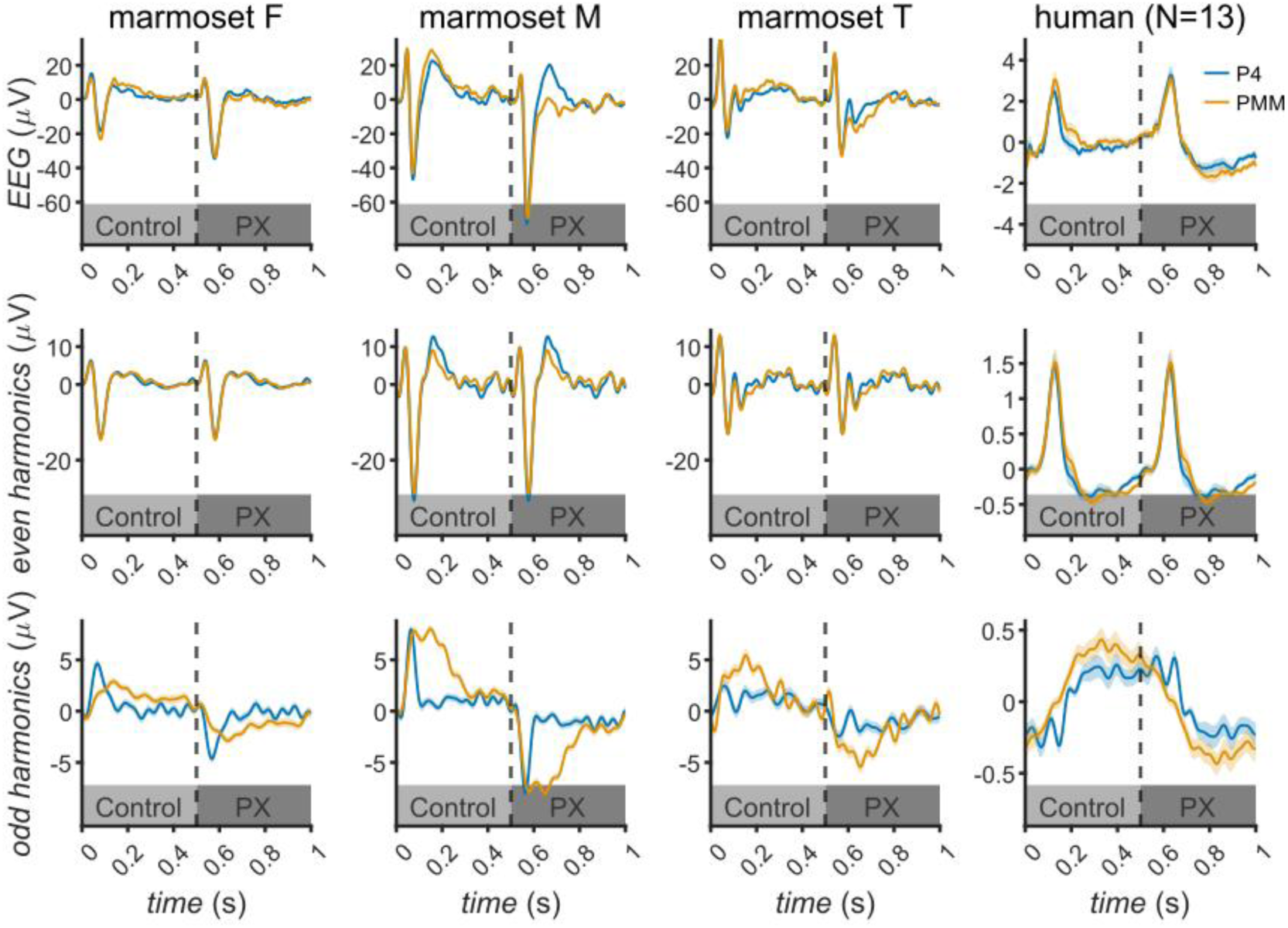
ROI-averaged EEG waveforms and harmonic reconstructions across marmosets and humans. Mean ± SEM signals are shown for three marmosets (F, M, and T) and a human group (N = 13) from an occipital ROI (marmosets: Oz, O3, O4; humans: six occipital electrodes). Columns correspond to subjects/groups and rows correspond to signal type. Top row: broadband time-domain EEG waveforms averaged across accepted stimulus cycles, spanning one full control–symmetry alternation cycle (dashed line indicates the control–symmetry transition at 500 ms). Middle row: time-domain reconstructions retaining only the even harmonics (2, 4, 6, … 20 Hz) of the 1 Hz stimulation frequency, reflecting transient responses to each image alternation. Bottom row: time-domain reconstructions retaining only the odd harmonics (1, 3, 5, … 19 Hz). In all panels, P4 (blue) and PMM (red) conditions are overlaid. Shaded regions indicate ± SEM across trials (marmosets) or across subjects (humans).

**Figure 4.**
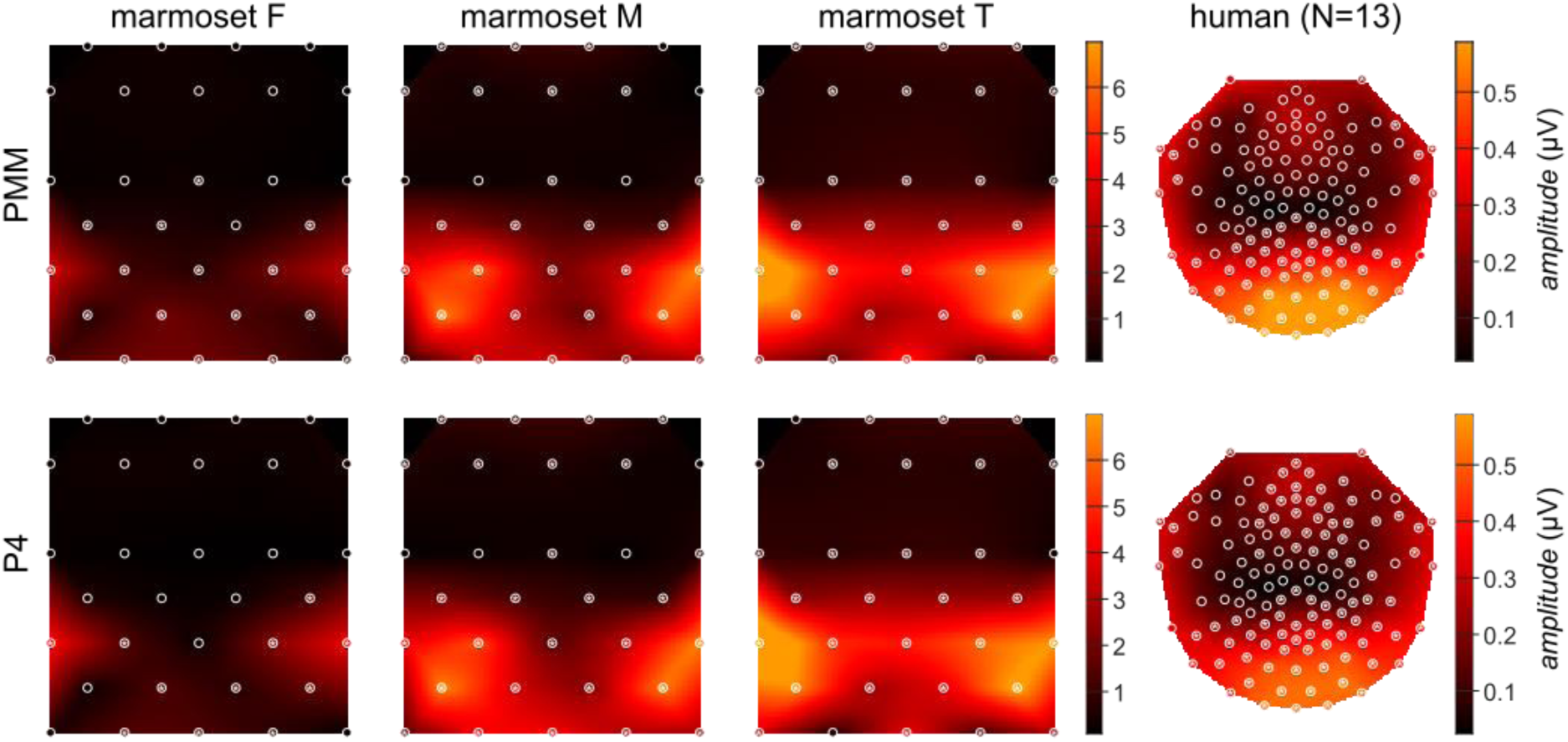
Topographic organization of even-harmonic steady-state EEG responses. Topographic maps show steady-state EEG response amplitudes at the first even harmonic (2f₁), plotted separately for each marmoset (columns: F, M, T) and for the human group (rightmost column). Rows correspond to the PMM condition (top) and the P4 condition (bottom). For both rows, colour indicates response amplitude: for marmosets, the magnitude of the mean complex Fourier coefficient at each electrode; for humans, the vector-projected mean amplitude across participants. Statistical significance in marmosets was assessed using the one-sample circular T² statistic [20] against the null hypothesis of zero mean complex response within each condition. For humans, significance was assessed using Hotelling’s T² test on the real and imaginary parts of the complex coefficients. Asterisks indicate electrodes showing statistically significant responses after false-discovery-rate correction (α = 0.01). White dots indicate electrode locations.

**Figure 5.**
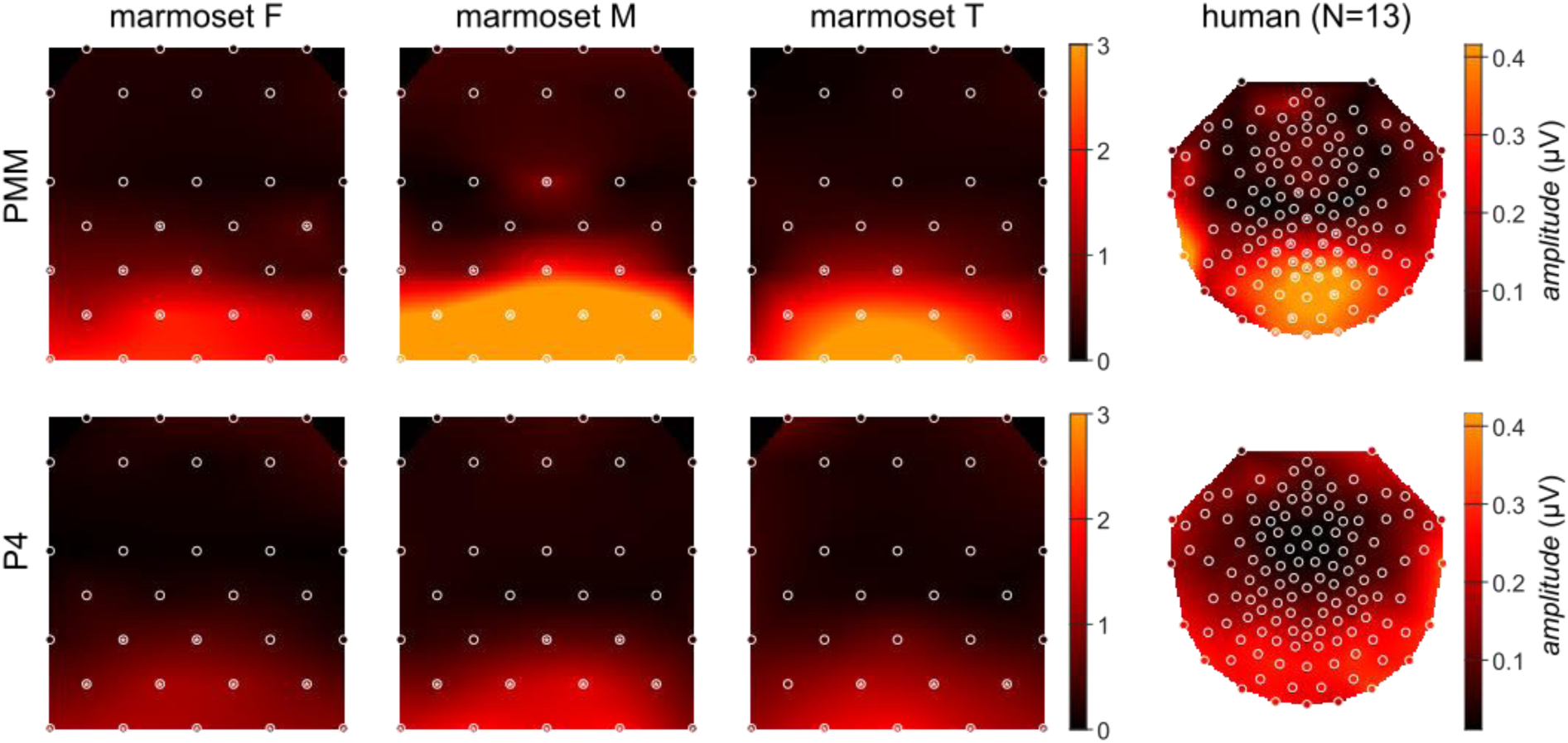
Topographic organization of odd-harmonic steady-state EEG responses. Same format as Figure 4, but for the first odd harmonic (**1f₁**). We next examined the first odd harmonic (1f), which captures the symmetry-related component of the response that differs between the two phases of the alternation cycle. In all three animals, 1f responses for both PMM and P4 were concentrated over posterior electrodes, with maxima centred on occipital sites (Figure 5). One-sample circular T² tests identified significant posterior odd-harmonic responses in each marmoset after false-discovery-rate correction (q < .01). Across significant marmoset electrode-by-animal observations, circular T² values ranged from 0.020 to 1.189 (p_FDR_ ≤ .008). In the human group, one-sample Hotelling’s T² tests identified significant posterior PMM responses after correction (T² (2, 11) = 26.65–70.81, p_FDR_ ≤ .010, Mahalanobis D = 1.432–2.334). These results indicate that symmetry-locked SSVEPs were robust at the single-sensor level across marmosets and the human group. Consistent with the human ROI-level effect, the first odd harmonic revealed systematic and spatially organized condition-related differences over occipital cortex.

To assess the overall strength of the odd harmonic symmetry responses relative to the image update response driven even harmonic response, we computed the root-mean-square (RMS) of the reconstructed odd and even harmonic waveforms (**Figure 6A**). We then divided the odd harmonic RMS for each wallpaper group, by the averaged even harmonic RMS over the two wallpaper groups. The results were consistent with individual harmonic analyses: Two out of 3 marmosets produced weaker odd harmonic RMS ratios for P4 than for PMM, but there was no consistent difference in response strength for the third marmoset, or for the human group (**Figure 6B**). Generally, the odd harmonic RMS ratios for marmosets were aligned in strength with the weaker end of the human distribution.

**Figure 6.**
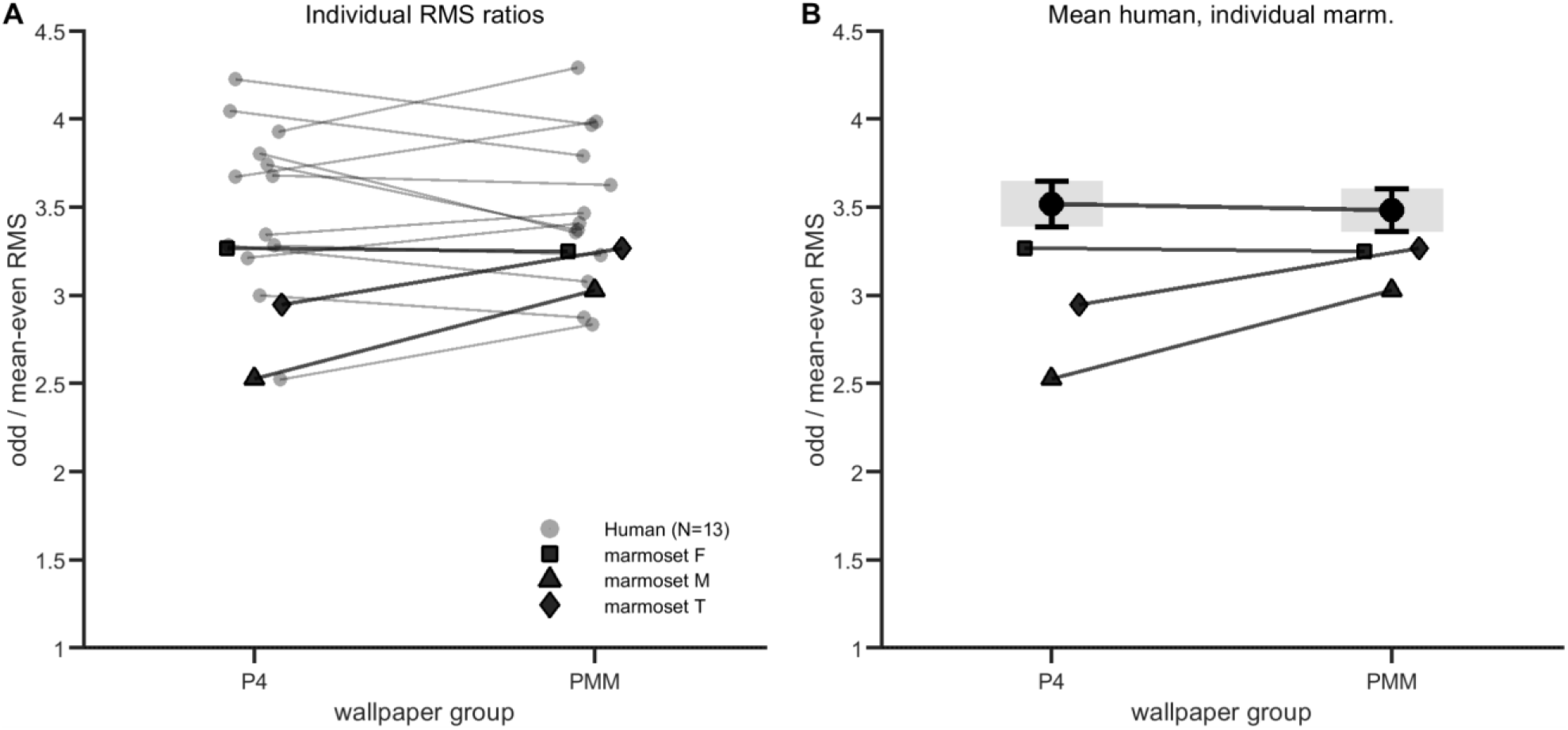
Odd-to-even harmonic response ratios in marmoset and human EEG. (A) Individual ratios of odd-harmonic RMS to mean even-harmonic RMS for P4 and PMM, shown for each human participant and each marmoset. (B) The same data shown as human group mean ± SEM with individual marmoset values overlaid. Ratios were computed from responses averaged across an occipital ROI (marmosets: O3, Oz, O4; humans: channels 70, 74, 75, 81, 82, and 83), using waveforms reconstructed from odd harmonics (1–19 Hz) and individual even harmonics (2–20 Hz).

## 4. Discussion

We used SSVEP to ask whether marmosets share the symmetry-processing computations that have been characterized in humans, and whether sensitivity to different symmetry types is similarly conserved. Skull-mounted EEG from three awake marmosets revealed robust, symmetry-specific responses over posterior cortex for both reflection (PMM) and four-fold rotation (P4). Reflection responses were comparable to those obtained from human participants viewing the same stimuli in relative amplitude, topography, and temporal profile. Rotation responses were also reliably present but transient, lacking the sustained late component that dominates the human response. This dissociation, obtained within the same animals and recording sessions, identifies the early feedforward computations underlying symmetry detection as candidate primate-general computations supported by conserved extrastriate mechanisms. The sustained processing that elaborates symmetry representations in humans—plausibly reflecting recurrent or feedback contributions—appears reduced in marmosets, and most markedly for rotation symmetry. These results establish marmosets as a tractable model for the conserved feedforward component of mid-level visual processing and motivate targeted investigation of the sustained processing that diverges between species.

### 4.1 Reflection symmetry as a conserved primate computation

The marmoset PMM response reproduces the defining features of the human symmetry SSVEP. Symmetry-specific signals were isolated in the odd harmonics of the stimulation frequency, segregated from transient image-update responses in the even harmonics—the same spectral dissociation that has been validated in human studies [12,13,18]. The topography was posterior and bilateral, consistent with extrastriate generators. Additionally, the response magnitude for PMM was comparable to that obtained in our human cohort using identical stimuli. The odd-harmonic waveform for PMM was sustained through each half-cycle in both species, indicating that the temporal profile of the reflection response is conserved across the marmoset-human divergence. Together, these features indicate that the marmoset posterior cortex performs an integration over reflection structure that operates on the same timescale, in the same spectral bands, and over a homologous cortical area as in humans.

This convergence aligns with the ecological logic of reflection symmetry. Reflection exists widely in bilateral symmetry in conspecifics, predators, and prey; and is a central signal in face, body, and animacy perception [21]. Both are domains where rapid, automatic detection carries clear adaptive value across primates. The presence of human-comparable reflection responses in marmosets— separated from the human lineage by around 40 million years [22]—suggests the underlying computation predates the platyrrhine–catarrhine divergence. This is consistent with macaque fMRI evidence of a homologous extrastriate network for symmetry [2] and extends that conservation to the temporal and spectral signatures accessible through EEG measurements of SSVEPs.

### 4.2 A transient–sustained dissociation for rotation symmetry

Reflection has been found to elicit larger responses than rotation within human visual cortex [11,12], and the same difference is present in macaque V3 and V4, where fMRI responses to reflection are approximately one third larger than to rotation [2]. In our current data, the human participants present a more subtle version of this effect. This may be due to the specific relatively low spatial frequency versions of the wallpaper patterns used here-prior work has shown that responses to reflection and rotation symmetries in wallpapers are differentially affected by spatial frequency content [23]. Two out of three of our marmosets had consistently larger responses for reflection than rotation, but in addition, all three demonstrated a remarkable difference in temporal profile: while they produced sustained odd-harmonic waveforms for PMM, their P4 responses were transient, returning to baseline much sooner than PMM. In our human data, both PMM and P4 produced reliably sustained responses, consistent with prior results [13].

Could these lack of sustained P4 responses be due to differences in gaze patterns between marmosets and humans? We consider this unlikely because the large size and tiled structure of the wallpaper patterns used here, means that the symmetry structure was present in the foveal region regardless of gaze location. Perhaps more importantly, any influence of eye movements would presumably apply to both reflection and rotation equally, rather than the rotation-specific difference between marmosets and humans we observe. The difference in temporal profile for rotation therefore appears to be a species-level effect layered on top of a within-species gradient that all three primates share. The fact that this lack of sustained responses is specific to rotation symmetry suggests that it is not a result of behavioural or other differences between marmosets and humans, but rather a species-level difference in cortical processing, layered on top of a within-species gradient that all primates share.

Two non-exclusive reasons could explain the species difference. First, rotation symmetry processing is more experience-dependent than reflection processing. Reflection symmetry is ubiquitous in the natural environment of any mobile primate, present in conspecifics, predators, and prey. Four-fold rotation symmetry, by contrast, is relatively rare in biological forms but on the other hand common in human-made artifacts such as tiles, mandalas, and mechanical parts—to which marmosets have negligible exposure. If sensitivity to higher-order rotation symmetry is partly shaped by visual experience with such patterns, weaker marmoset responses would follow. Second, humans and marmosets may differ in the cortical dynamics that follow initial symmetry detection. In humans, the odd-harmonic response to both reflection and rotation persisted throughout each half-cycle. In marmosets, the reflection response was similarly sustained, whereas the rotation response was clear and reliable but transient, returning to baseline well within the half-cycle. This shows that in our paradigm, marmosets do exhibit robust responses to rotation symmetry, but with more transient temporal dynamics, compared to humans. A transient response is most readily explained by feedforward processing, whereas a sustained response may be the result of additional recurrent or feedback activity that further elaborates the symmetry representation. This fits comparative connectomics showing that interareal projections in marmoset cortex are, on average, more feedforward in laminar origin than in macaque [17]. It is an open question why the species difference emerges for rotation and not reflection—we may speculate that the sustained component of the response demands more extensive feedback connectivity for rotation than reflection. Laminar recordings in marmoset extrastriate cortex will shed light on the feedforward and feedback contributions to each response component and help resolve this question.

Our findings cannot adjudicate between these accounts, and they are not mutually exclusive. Distinguishing an experience-dependent process from cortical specialization will require developmental manipulations or comparisons across rearing environments. What our results establish, is that the rotation-specific absence of sustained responses is robust within the same animals and recording setup and does not reduce to a low-level or non-specific difference.

Two features of the data support this. First, the even-harmonic responses that index the transient reaction to each image update were closely matched between PMM and P4 in both waveform and scalp topography. This indicates that the two wallpaper groups were equated for low-level stimulus drive and that the divergence was confined to the odd, symmetry-specific harmonics. Second, odd- and even-harmonic amplitudes were uncorrelated on a trial-by-trial basis, and the condition effect at the first odd harmonic survived partialling out even-harmonic amplitude. This rules out a global gain change or non-specific signal-to-noise difference as its source.

The even-harmonic condition differences, though modest and inconsistent in direction, suggest that the image-update response was subject to top-down gain modulation that differed between pattern types. One candidate for this is differential attention or arousal to reflection versus rotation [18].

The subject-specific direction of the effect is consistent with idiosyncratic engagement rather than a stimulus confound. Importantly, this modulation cannot explain the symmetry results: the odd-harmonic PMM–P4 difference was statistically independent of even-harmonic amplitude on a trial-by-trial basis, so the symmetry-specific signal is dissociated from any attentional gain change.

### 4.3 Cortical generators

The spatial distribution of symmetry-specific SSVEPs provides initial constraints on their likely cortical sources. Anatomical and functional mapping studies indicate that marmoset V1 occupies the occipital pole, V2 forms an anterior belt, and dorsal extrastriate areas —including partial homologies to human regions V3 and V6—respectively lie laterally and medially on smooth cortical surface, in contrast to macaques where these regions are buried in sulci [15,16,24–26]. Our grid covers much of the dorsal surface of the skull, with the O row placed over the occipital pole, the PO row near the V1–V2 border, and the P row over dorsal extrastriate cortex encompassing V2 and V6. Symmetry-specific odd-harmonic responses peaked in the O row but remained robust at PO and P, consistent with strong symmetry-locked activity in posterior cortex with contributions from more dorsal sites.

These findings align with human and macaque imaging. Human fMRI shows symmetry responses emerging in V3 and strengthening along the ventral stream through V4, VO1, and lateral occipital cortex [1,13,27]. In macaques, fMRI reveals a comparable network spanning V2, V3, V4/V4A, and PITd with tuning properties closely matching those in humans [2]. Skull-recorded EEG does not allow for localizing the symmetry-related SSVEP unequivocally to specific areas, but the posterior focus and O–PO–P gradient are consistent with generators in early visual cortex and dorsal extrastriate regions, providing guidance to future single-unit recordings.

### 4.4 Advantages of novel skull-mounted EEG

A key methodological contribution of this study is a chronic, flexible EEG grid that mounts directly on the marmoset skull. The grid covers most dorsal cortex from prefrontal to occipital areas, and is thus suitable for studying diverse cognitive processes. We used it head-fixed here, but it also supports head-free recording in a chair, and wireless recording when paired with a compatible head stage.

The grid offers three advantages over existing approaches. First, scalp EEG requires daily electrodes application [28–31]. Our skull-mounted interface is mechanically stable, supporting consistent recordings across months and is plug-and-play on a daily basis. This preserves animal motivation and alertness for the task. Second, compared to ECoG grids which require surgical removal and replacement of large sections of skull [29], our approach avoids this and minimizes tissue damage and infection risk, and remains compatible with later invasive procedures. Laminar probes or optical windows can be inserted through separate craniotomies, allowing future experiments to directly relate SSVEP measures of symmetry processing to local spiking and field activity in targeted visual areas. Third, we designed the posterior montage and electrode spacing to mirror human occipital SSVEP arrangements. This yields responses that are spatially and spectrally comparable to human EEG, strengthening the translational link between marmoset and human studies of mid-level form processing [13,15,18]. The grid is also safe for magnetic resonance scanners and pairs naturally with fMRI and with head-free eye-tracking and receptive-field mapping [32], supporting studies of naturalistic vision and behaviour.

### 4.5 Limitation and future directions

Two limitations bound the current conclusions. First, our stimulus set included only two wallpaper groups. Extending the current paradigm to other wallpaper groups—including those with higher rotation orders and glide-reflection symmetry—will determine whether marmoset symmetry representations recapitulate the detailed hierarchical structure observed in human SSVEPs and psychometric thresholds [12,13]. Such an extension would also clarify whether the reduced P4 response reflects a specific deficit for high-order rotation or rotational symmetry in general. Second, our passive-viewing design does not support inferences about active discrimination performance or to test whether task plays a role in marmoset symmetry responses. Future work combining SSVEP with behavioural readouts of symmetry detection in marmosets will help link the neural signal to perceptual and cognitive consequences

### 4.6 Conclusion

This study provides the first electrophysiological measurement of symmetry processing in marmosets and reveals a clear dissociation: Reflection symmetry is processed with human-like robustness and temporal dynamics, while rotation symmetry produced a reliable but transient response that lacks the sustained late component seen in humans. The shared reflection signature— well-matched in spectral structure, topography, and magnitude to human SSVEP—identifies reflection-symmetry detection as a candidate primate-general computation supported by conserved extrastriate mechanisms. The transient rotation-symmetry difference, by contrast, points to a specific dissociation in late-stage symmetry processing between species, plausibly reflecting differences in recurrent or feedback contributions that elaborate the initial feedforward representation. Combined with the marmoset’s accessible cortex and growing experimental toolkit [15,33–35], the human-comparable symmetry SSVEP positions this species as a tractable platform for mechanistic dissection of the conserved component of mid-level vision. Our findings also motivate targeted investigation of the recurrent and feedback contributions to symmetry processing across species. More broadly, this gives perceptual-domain evidence for a cross-species difference in feedback processing that has so far been described mainly based anatomy and in executive functions, and it shows how a human-comparable EEG signal in a primate model can pinpoint where in the processing hierarchy that difference emerges.

## Acknowledgement

We thank Shuyuan Lyu for performing MRI-based 3D skull reconstruction for 2 of the 3 subjects. We thank Christopher Giverin for his help with the custom design of the head chamber and surgical mask.

## Data availability

All data and analysis code are available via Zenodo (DOI: https://zenodo.org/records/22379945).

